# Investigation of Biomedical Cell Image Cryptography Based on RC4 Technique

**DOI:** 10.1101/2023.12.29.573618

**Authors:** Song Jiang, Ela Kumar

**Affiliations:** University of Houston, Texas, USA; Department of Computer Science and Engineering, KLEF, Vaddeswaram, India

**Keywords:** Biomedical, Image, RC4, Cryptography

## Abstract

Enormous data permeates various facets of social life, including public administration, community services, and scientific research, owing to the advent of new technologies. With the escalation of data transportation concerns, its significance in our daily lives has magnified. Particularly in the medical field, preserving information integrity during data transit has become an urgent priority. For instance, safeguarding massive electronic medical files like MRI images from cyberattacks and data loss poses a formidable challenge. Encryption and decryption techniques are employed as a solution. While numerous image types have undergone testing using cryptography techniques, biomedical images have received limited attention. In this article, We implemented RC4, pixel shuffling, and steganography to encrypt and decrypt cell images. Evaluating the encryption and decryption performance of these cell images involved testing cell viability, live/dead ratio, and morphology.

## 1. Introduction

Biomedical applications such as diagnostic imaging, tissue regeneration, and the use of antibacterial agents have seamlessly integrated into our daily lives. These applications effortlessly support various aspects of healthcare and well-being, becoming integral parts of our daily routines.[1-8] A multitude of these applications relies on image analysis, encompassing X-rays, MRIs, and cellular imaging.[9] Annually, these advancements benefit thousands of patients dealing with various conditions like headaches, cancer, and abdominal pain. The integration of imaging technology with other diagnostic methods aids in making preliminary diagnoses to determine and confirm the cause of ailments.[10] The evolution of medical imaging technology in clinical diagnosis holds tremendous importance, encompassing technology updates and technician training. The aging population and the prevalence of cancer among younger patients have accelerated the rapid development of medical imaging.[11] Leveraging digital imaging technology allows for more accurate and faster patient diagnoses, leading to earlier treatment interventions. Moreover, the utilization of machine learning and statistical analysis in medical imaging has become increasingly pivotal in clinical diagnoses, given its highly integrated, digitized, networked, fused, and standardized nature.[12-15] Clinical imaging diagnosis significantly contributes to enhancing diagnostic accuracy and reducing misdiagnoses, thereby improving patient care.[16]

With the continuous advancement of high technology, an increasing volume of information is transmitted through networks. Digital images have gained immense popularity among people due to their visibility, intuitiveness, and other characteristics, becoming a pivotal means of information exchange.[17] The transmission of digital images over the internet significantly impacts people’s lives. Despite the internet being an open and shared platform, transmitting image information via the internet entails the risk of information leakage.[18] Consequently, ensuring image security has become a crucial concern. Employing image encryption technology stands as the most effective method to address this issue, ensuring robust image security. However, discovering a secure and effective image encryption method remains a considerable challenge.

Image encryption technology primarily consists of two different types. One type involves encrypting an image into a ciphertext image. When the ciphertext image and the plaintext image obtained are of the same size, it is termed symmetric encryption. The other type compresses and encrypts the image information. This process results in a ciphertext image size smaller than that of the plaintext image, known as asymmetric encryption.[19]

## 2. Related Research

In the biomedical field, the encryption of images has reached a mature stage of development. There exist numerous encryption methods tailored for securing these images. Each method boasts its own unique advantages and disadvantages. For instance, Bhaskar Panna devised an encryption technique employing the fractional discrete cosine transform in combination with chaotic function. His method ensures the security of medical data by employing a chaotic map on the fractional discrete cosine transform coefficients of medical images, offering a high degree of freedom for their encryption.[20] Similarly, C. Priya developed a medical image encryption method based on a highly efficient Encryption-Compression system with lossless compression. This approach involves mathematical permutations to provide a reasonably high level of security. Moreover, it integrates run-length coding for efficiently compressing the encrypted images.[21]

Fascinatingly, Pang Ming devised an encryption method and conducted a security analysis of medical images using a stream cipher enhanced logical mapping. His approach relies on the Chebyshev map to generate the encryption key. A sequence of coding operations is implemented to establish the initial value before initiating image chaos processing. Ultimately, through integration with logical mapping, the original image information undergoes encryption through chaos from the X and Y dimensions.[22]

## 3. Motivation

Although many kinds of images have been investigated using various types of encryption and decryption techniques, few have focused on the biomedical area, particularly in the analysis of cell images. Here, we have selected cell images as the target for encryption and decryption, considering their fundamental role in biomedical research. We have opted for the RC4-pixel shuffling steganography system as the encryption and decryption method due to its strength in preventing attacks from hackers without causing information loss.

## 4. Methodology

### 4.1 Dataset

There are three types of cell images used in this article: cell proliferation, survival tests, and cell morphology. All images in this article were randomly selected online.

### 4.2 Methods

The methods for encryption and decryption of cell images are based on the RC4-pixel shuffling-steganography system. The flow chart of procedure shows in figure 2.

**Figure 1.**
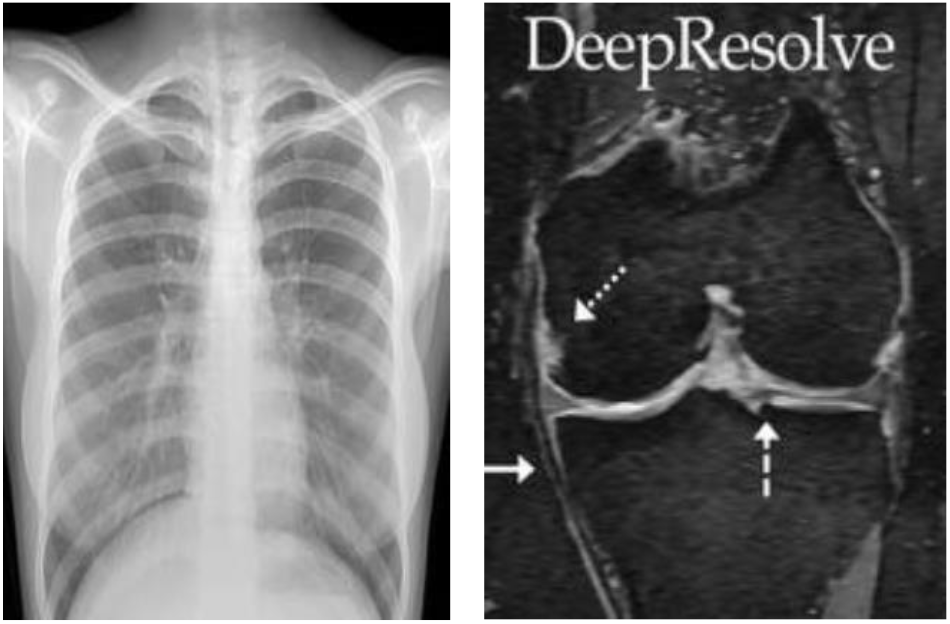
Typical Medical Images of X-Ray and MRI

**Figure 2.**
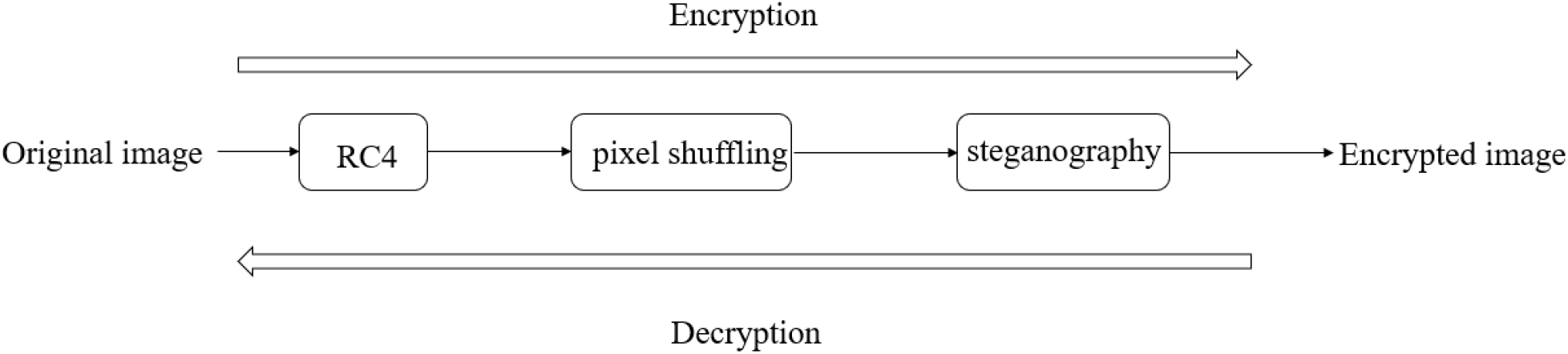
The flowchart of RC4-pixel shuffling steganography system

#### 4.2.1 RC4

RC4 is a stream cipher algorithm, which is described in the figure 3. It is a non-linear transformation-based algorithm commonly used in the data transportation. It includes two aspects: key schedule algorithm and pseudo random generation algorithm (PRGA). Key schedule algorithm use variable-length keys to perform random sorting to generate the initial state of the key stream. PRGA is used to refer to the initial state to generate the corresponding key stream sequence, and then generate the corresponding ciphertext by XORing with the plaintext. Here, The encryption principle of the RC4 algorithm shows in the figure 4. The first step is to initialize the two vector registers S and T according to the established algorithm. The second step is to rearrange the register S. The last step is to generate the corresponding The key stream k, followed by storing and participating in the subsequent plaintext encryption operation.[23]

**Figure 3.**
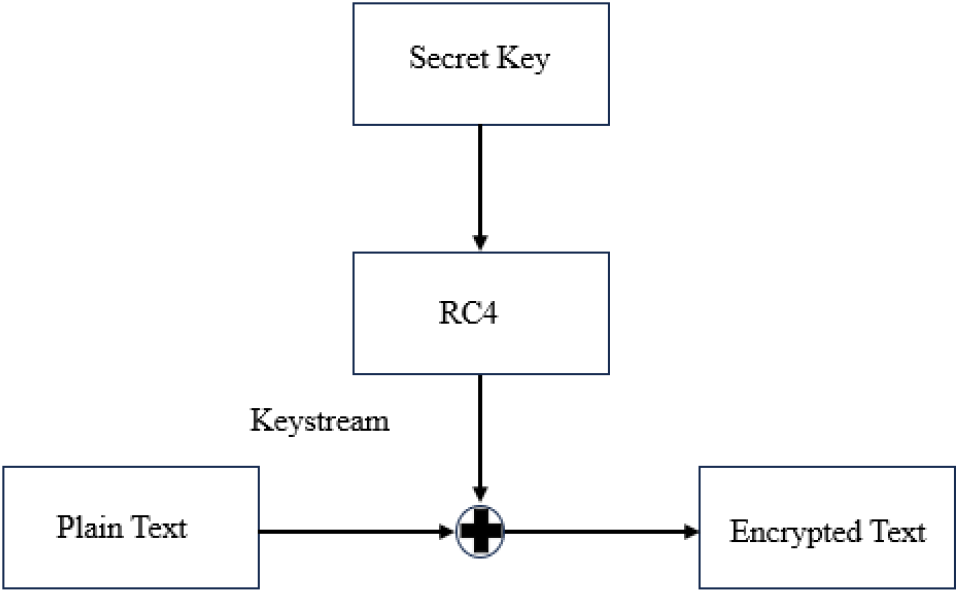
The flowchart of RC4

**Figure 4.**
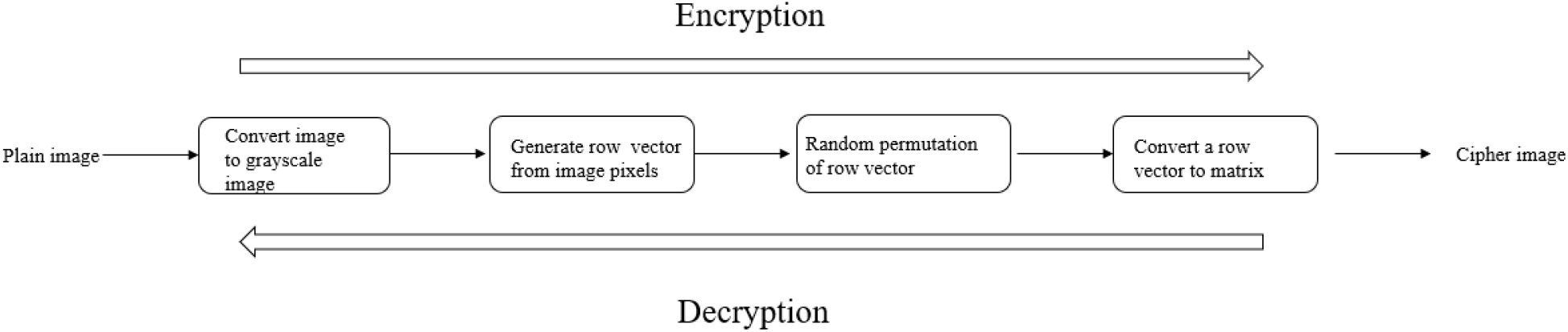
The flowchart of pixel shuffling

#### 4.2.2 Pixel Shuffling

After RC4 encryption, pixel shuffling is applied to further encrypt cell images. Pixel shuffling is used to permutate the position of the cell images. Figure 5 shows the procedure of pixel shuffling. First original cell image is imported and read, followed by converting the original cell image to grayscale one. Then, a row vector from image pixel is generated and randomly permutated, followed by converting a row vector to matrix.[24]

**Figure 5.**
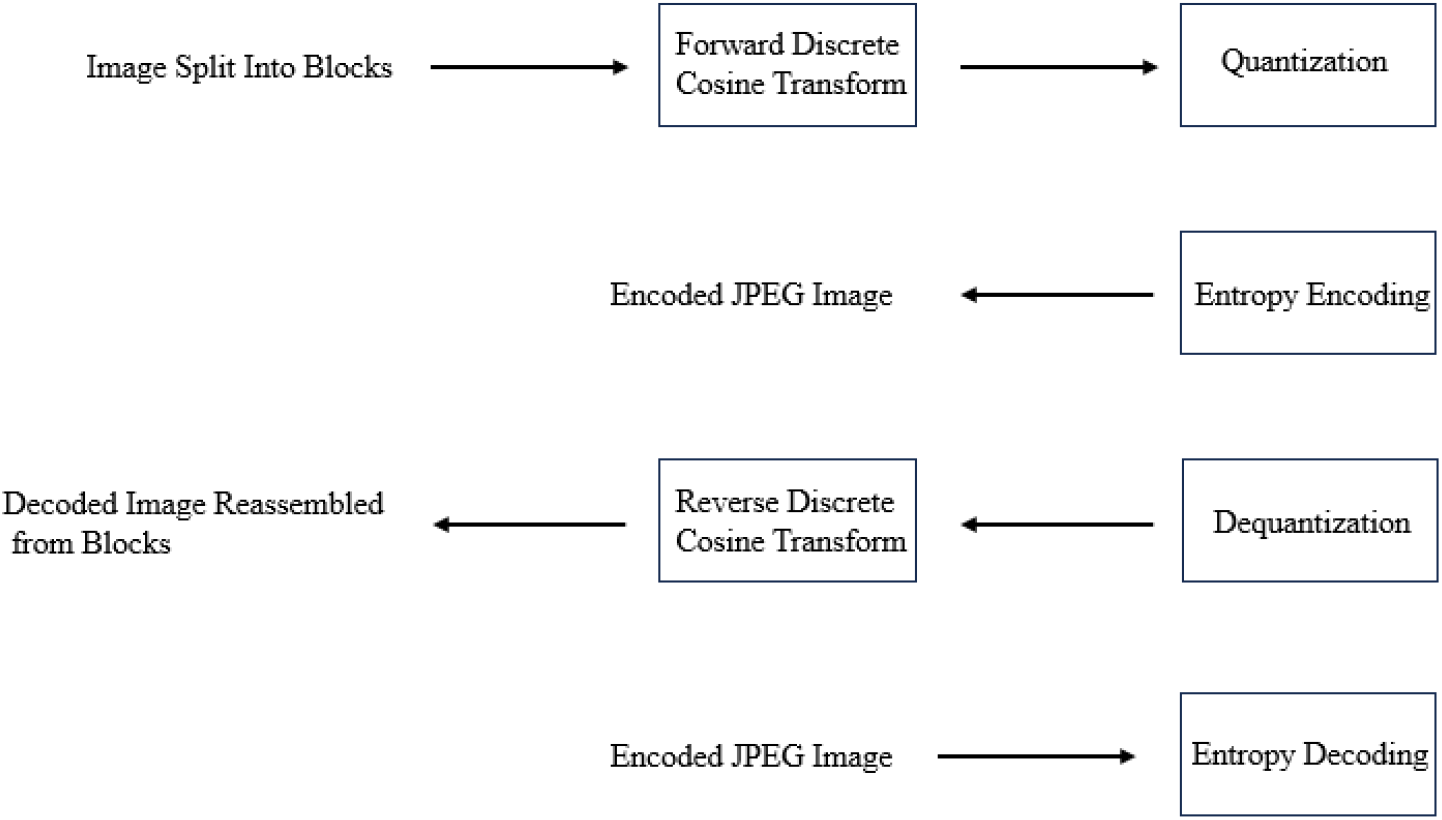
The flowchart of steganography

#### 4.2.3 Steganography

Steganography is used to hide the secret information in the public information medium for transmission. The model of the steganography system is shown in figure 8. The carrier used to hide the message is called the cover signal, which can be digital image, video, audio or even text data. Another input in the embedding process is the embedded object. It can be various forms of digital information represented by a bit stream. In order to improve confidentiality, sometimes the embedded object is encrypted. In the extraction process, it is divided into two types: non-blind extraction and blind extraction, according to whether the participation of the carrier object is required. The key generator generates the embedded key and extracts the key according to the security parameters.[25]

### 4.3 Measurement

The measurement is conducted by ImageJ, which is a specific software to analyze the encrypted and decrypted cell images.[26] The measurements include viability, survival ability and morphology. Viability is the number of cells. Survival ability is the ratio between number of live cells and that of dead cells. Morphology is the length of the cell and the height of the cell. The results of measurements of original cell images are compared with decrypted cell images.

## 5. EXPERIMENTS AND RESULTS

The figure 6 shows the sample original cell image, encrypted image and decrypted image. The encryption of cell image is processed by RC4, followed by pixel shuffling and steganography. The decryption of cell image is vice versa. From these figures, it is obvious that all images proves that the RC4-pixel shuffling-steganography system can be applied to encrypt and decrypt cell images successfully.

**Figure 6.**
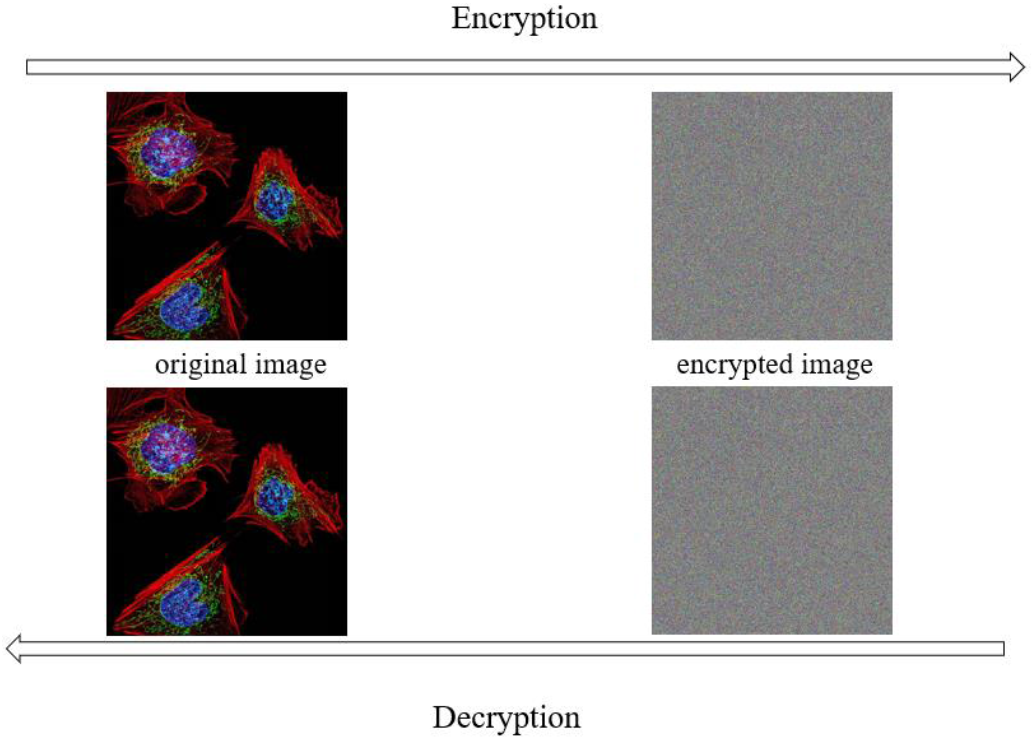
The sample cell images of encryption and decryption

### 5.1 Comparison

#### 5.1.1 Viability

Figure 7 illustrates the cell proliferation, with the cell count determined using ImageJ. In the original image, the cell count is 356. Similarly, the cell count in the decrypted image is also 356.

**Figure 7.**
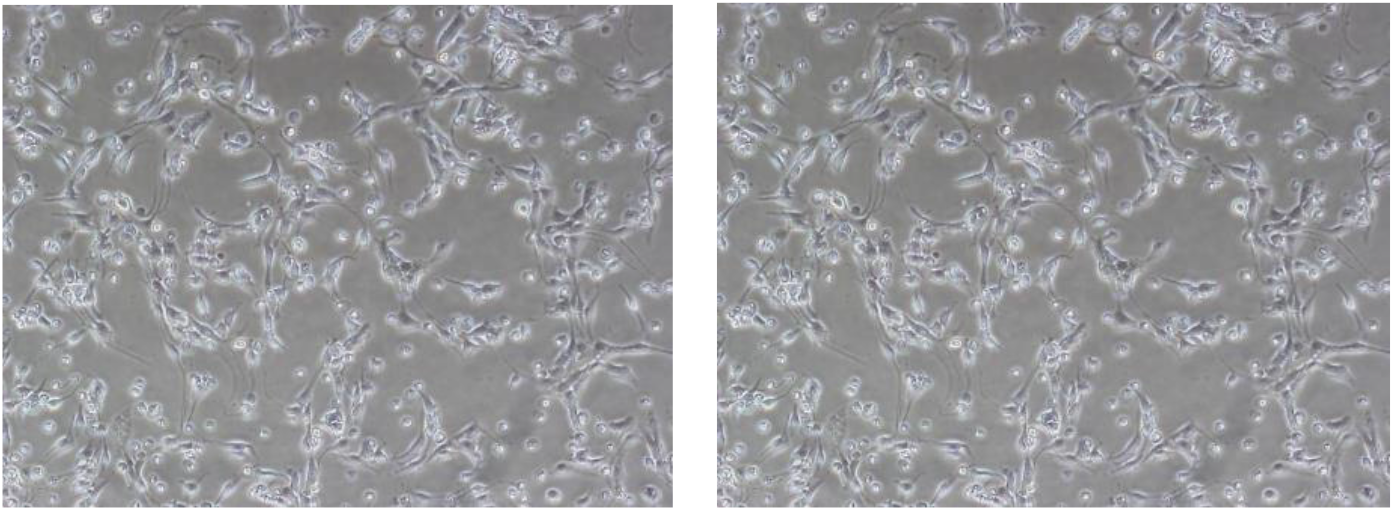
Cell proliferation of original cell image (left) and decrypted image (right)

**Figure 8.**
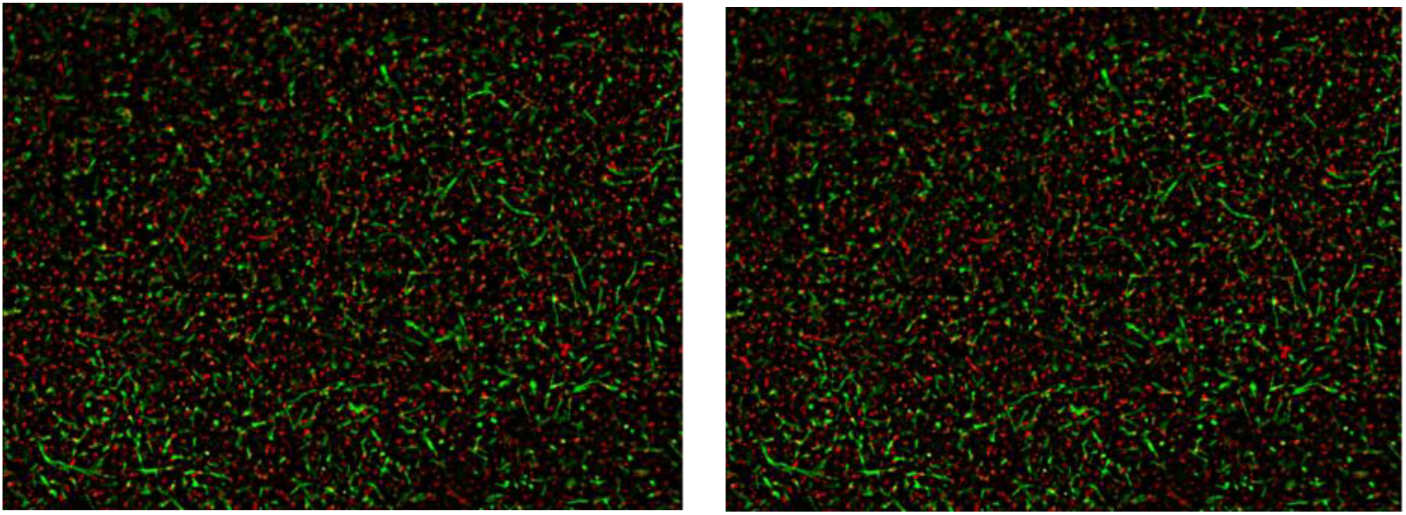
Cell survival ability of original cell image (left) and decrypted image (right)

#### 5.1.2 Survival Ability

Figure 8 displays the survival ability of cells, where red spots represent dyed dead cells and green spots indicate live cells. The survival ability is determined by calculating the ratio between dead and live cells. The survival ability, based on the original cell image, is 65.1%. Similarly, the survival ability derived from the decrypted cell image is also 65.1%.

#### 5.1.3 Cell Morphology

Figure 9 displays the cell morphology of both the original and decrypted images. Two parameters were measured: the length and height of the cell. In the original image, the cell’s length measures 14 µm, and its height measures 9 µm. Similarly, in the decrypted image, the length and height of the cell also measure 14 µm and 9 µm, respectively.

**Figure 9.**
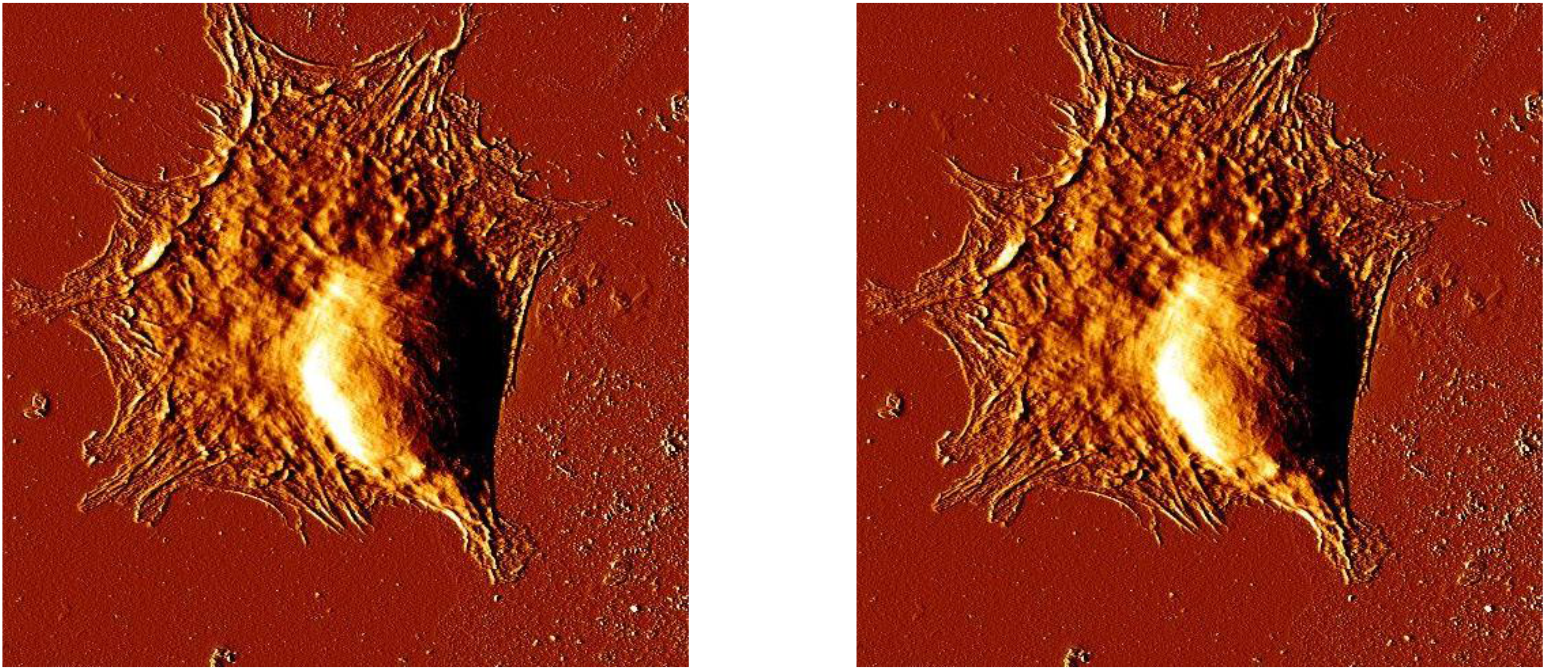
Cell morphology of original cell image (left) and decrypted image (right)

## 6. Conclusion

Based on the aforementioned analyzed results regarding cell image viability, survival ability, and morphology, it is evident that all the test results are identical between the original image and the decrypted image. Consequently, it can be concluded that cell images can be encrypted and decrypted using RC4, pixel shuffling, and steganography without any loss of information. Therefore, cell images can be safely transported.

## 7. In the Future

Due to time constraints, several tasks remain to be completed to perfect this project. For instance, the RC4-pixel shuffling steganography system could be optimized. Furthermore, additional tests comparing the original cell image with the decrypted image can be conducted.

